# Lizards on a sky archipelago: Genomic approaches to the evolution of the mountain genus *Iberolacerta*

**DOI:** 10.1101/2025.06.04.657811

**Authors:** Adrián Talavera, Bernat Burriel-Carranza, Gabriel Mochales-Riaño, Maria Estarellas, Oscar Arribas, Héctor Tejero-Cicuéndez, Judit Salces-Ortiz, Daniel Fernández-Guiberteau, Fèlix Amat, Manel Niell, Rosa Fernández, Alexander S. Mikheyev, Salvador Carranza

## Abstract

The mountain-dwelling lizards of the genus *Iberolacerta* inhabit several isolated massifs across central and southwestern Europe. Their restricted and fragmented ranges, coupled with high altitude specialization in most species, entail a significant threat in the context of climate change for this group of lizards that has attracted interest from different fields. On the one hand, the alpine confinement of these relict species precedes the Pleistocene glacial cycles, and a few hypotheses have been proposed to explain it: from competitive exclusion by the wall lizards of the genus *Podarcis*, to adaptations to either cold or hypoxia, that would prevent them from expanding into lowlands. On the other hand, extensive research on chromosome evolution has shown *Iberolacerta* karyotypes to fairly differ from other lacertid lizards, exhibiting reductions in chromosome numbers and multiple sex chromosome determination systems. Here we present a chromosome-level genome assembly for *Iberolacerta aurelioi*, an Endangered rock lizard endemic to the Pyrenees. This genome has shed light on a genome architecture shaped by chromosome fusions, whose adaptive potential we discuss, as well as on expression shifts towards a hemoglobin isoform of enhanced oxygen affinity, as an adaptation to altitudinal hypoxia. In addition, medium coverage whole-genome sequencing data from 12 representatives encompassing all species and subspecies within the genus allowed us to address phylogenomic relationships, unveiling introgression events, gathering evidence in support of the competitive exclusion hypothesis through past demographic inference, and providing insights into homozygosity burdens, which offer valuable information for conservation efforts.

## Introduction

Southern Europe stands as a diversity hotspot for Lacertidae lizards, where progressive global cooling over the last 10 million years triggered the expansion of their bioclimatic niches (Garcia-Porta et al., 2019). Mostly restricted to highlands in southwestern Europe, we find an assemblage of old lacertid species: the rock lizards of the genus *Iberolacerta* (Fig. 1ABC). This genus has been subject of thorough phylogenetic research in the last decades (Arribas et al., 2006, 2014; Arribas & Carranza, 2004; Carranza et al., 2004; Crochet et al., 2004; Mayer & Arribas, 2003, 1996; Mouret et al., 2011; Remón et al., 2013), settling to the existence of eight species: one species, *Iberolacerta horvathi*, inhabiting the Eastern Alps and Northern Dinaric Ranges; the so-called *Pyrenesaura* subgenus found in the Central Pyrenees, composed by *Iberolacerta bonnali*, *Iberolacerta aranica* and *Iberolacerta aurelioi*; and four species with different subspecies occurring in Central and Northwestern Iberia. Among the latter, *Iberolacerta monticola monticola* is the most widespread taxon, occurring in the Cantabrian Mountains, and in several scattered lowland relict populations, which even reach near-sea elevations in deep fluvial gorges (Remón et al., 2013). Additionally, an isolated *I. m. monticola* population is present in the westernmost Central System, in the Serra da Estrela (Portugal). South to the Cantabrian Mountains, the León Mountains are inhabited by *I. monticola astur* and the closely related *Iberolacerta galani* (Arribas et al., 2006, 2014). Finally, further east in the Central System beyond *I. monticola monticola*’s range in Serra da Estrela, we find the Endangered microendemic *Iberolacerta martinezricai*, restricted to the Sierra de Francia Massif, and the more widespread *Iberolacerta cyreni*. The latter is represented by the subspecies *I. cyreni castiliana*, found in Béjar, Gredos, and smaller neighboring ranges, and *I. cyreni cyreni*, which inhabits the Guadarrama mountain range (Fig. 1).

**Fig. 1.**
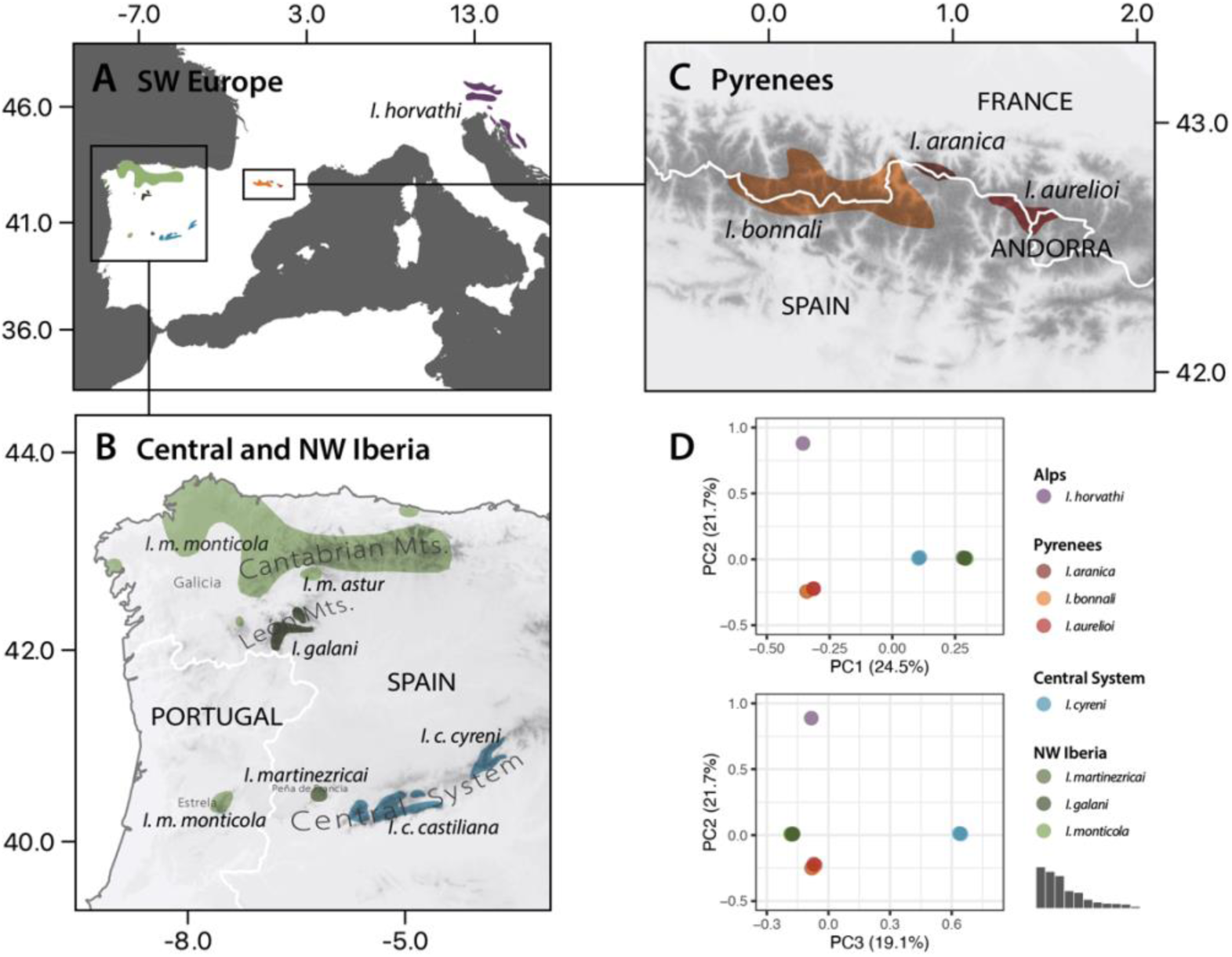
Distribution of the rock lizards of the genus *Iberolacerta*, endemic to southwestern Europe, and their genomic clustering into four clades. A-C) Ranges of all *Iberolacerta* species, mainly restricted to montane or alpine areas. Darker greys depict increasing altitudes. *Iberolacerta horvathi* inhabits the eastern Alps and northern Dinaric Alps, being the only species in the genus not inhabiting the Iberian Peninsula. Four species inhabit several ranges in Central and northwestern Iberia (B). The Central System is inhabited by *I. cyreni*, *I. martinezricai* and a southern isolated population of *I. monticola monticola*, from East to West. Other populations of *I. m. monticola* occupy the Cantabrian Mountains and some coastal populations in Galicia and Asturias (Spain). *Iberolacerta galani* and the subspecies *I. monticola astur* inhabit the León Mountains, south to the Cantabrian Mountains. Three more species, comprising the subgenus *Pyrenesaura*, inhabit the Central Pyrenees: *I. bonnali*, *I. aranica* and *I. aurelioi*, from West to East (C). D) Genomic PCA with >125k uSNPs, splitting the genus into four main clades: Alps, Pyrenees, Central System (East) and NW Iberia. Relative explained variance by PC at the bottom right corner.

Although all *Iberolacerta* species are characterized by their montane preferences (Carranza et al., 2004), with some exhibiting the lowest habitat diversity among Western European lacertids (Escoriza et al., 2023), the three species of the subgenus *Pyrenesaura* particularly stand out as strictly high-mountain dwellers, inhabiting the Pyrenean massifs at elevations ranging from approximately 1,900 to over 3,000 meters above sea level (Arribas & Galán, 2005). This alpine specialization results in strong population isolation. Similarly, other non-*Pyrenesaura* species such as *I. martinezricai* also display extreme habitat restriction, being confined to a 12–15 km² area in the Spanish Central System (Carbonero et al., 2016). Small and isolated populations like these are especially vulnerable to extinction due to demographic and environmental stochasticity, as well as genetic threats such as inbreeding (Remón et al., 2012). As a result, two *Iberolacerta* species — *I. martinezricai* and *I. aurelioi* — are currently listed as Endangered (Bowles, 2024a, 2024b).

Apart from high altitude specialization, all these rock lizard species share old origins, indicating long persistence in mountains preceding —and despite— the Pleistocene glacial cycles (Carranza et al., 2004; Crochet et al., 2004; Mouret et al., 2011). This long alpine confinement has been hypothesized to result from the simultaneous spread of wall lizards (*Podarcis*) and the subsequent competitive exclusion of *Iberolacerta* species (Carranza et al., 2004). However, contrasting evidence regarding competition between these genera has been reported among species (Galán, 2019; Monasterio et al., 2010; Žagar et al., 2015). Alternatively, adaptation to cold conditions has been proposed as another explanation for *Iberolacerta*’s alpine confinement, backed up by evidence on thermal constraints for embryonic development (Monasterio et al., 2011, 2016) and high metabolic performance, which can be too costly to maintain at higher temperatures (Žagar et al., 2018). Regardless of whether cold adaptations or *Podarcis* competitive exclusion confined *Iberolacerta* to highlands, climate change constitutes a significant threat. While mountains may experience greater warming rates than the global average (Rangwala & Miller, 2012), and high-altitude ectotherms are more sensitive to increasing temperatures (Feldmeier et al., 2020; Kingsolver et al., 2013; Paaijmans et al., 2013), *Podarcis* is expected to expand its range upslope leveraging global warming, which might foster potential competition or expose *Iberolacerta* to novel diseases (Gangloff et al., 2021; Ortega et al., 2016b).

Furthermore, a third, non-exclusive, and recently proposed hypothesis to explain alpine confinement is that *Iberolacerta* has adapted to high-elevation hypoxia to a point where relative “hyperoxia” at lower elevations renders organismal malfunction (Gangloff et al., 2021). Although the physiological effects of hyperoxia are not well studied, experiments on the Pyrenean *I. bonnali* have supported this hypothesis (Gangloff et al., 2021). Similar studies of longer duration have reported plastic physiological responses to hyperoxia in terms of hemoglobin (Hb) concentration falls in *I. cyreni* (Megía-Palma et al., 2020). This suggests that *Iberolacerta* may possess Hb of increased oxygen affinity, in line with the hyperoxia-as-constraint hypothesis and making *Iberolacerta* species good models to study adaptation to low oxygen availability.

Moreover, *Iberolacerta* species have garnered attention not only from the biogeographic and ecological points of view, but from the uniqueness of their karyotypes as well (Arribas & Odierna, 2004; Naveira et al., 2023; Odierna et al., 1996; Olmo et al., 2004; Rojo et al., 2013). Extensive research on chromosome evolution has revealed a large variation in diploid number among —and even within— species: from 2n= 36 of NW Iberian species to 2n=22 in some *Pyrenesaura* populations, all lacking the ancestral lacertid microchromosome pair (Naveira et al., 2023; Odierna et al., 1996), as well as noteworthy diversity in the differentiation of sex chromosomes, from cryptic to highly heteromorphic ZW chromosomes and even multiple Z_1_Z_2_W systems (Odierna et al., 1996; Rojo et al., 2013). Unfortunately, the lack of fully assembled genomes has hindered a more thorough understanding of the chromosomal arrangements that shaped such unprecedented diversity.

Here, we present, within the framework of the Catalan Initiative for the Earth Biogenome Project (Corominas et al., 2024), a chromosome-level *de novo* genome assembly for the Aurelio’s rock lizard *Iberolacerta aurelioi* —an Endangered species endemic to the Pyrenees (Pérez-Mellado et al., 2009)— in which we have explored chromosomal rearrangements and selection on hemoglobin-encoding genes as an adaptation to hypoxia. In addition, we have sequenced medium-coverage whole genomes from 12 representatives of all known *Iberolacerta* lineages to infer phylogenomic relationships, introgression events, and demographic oscillations, and to estimate inbreeding through runs of homozygosity. Our results confirm the old origins of most of species but reveal the recent divergence of *I. martinezricai* and *I. galani* during the Pleistocene, coeval to the subspecific diversification of other *Iberolacerta* species. Population expansions and contractions during this period are reported for NW Iberian and Central System taxa, in agreement with the occurrence of introgression within the first clade. In contrast, the relict character of Alpine and Pyrenean species dates back to the Pliocene, with bottlenecks matching *Podarcis* diversification and reflected in longer runs of homozygosity and lower heterozygosity. Finally, *I. aurelioi*’s genome sheds light on seven chromosomal fusions and an expression shift towards a former-minor hemoglobin isoform with enhanced oxygen affinity.

## Materials and methods

### DNA extraction, library preparation and sequencing

We extracted genomic DNA from tail tips for 12 specimens representing all taxa within the genus *Iberolacerta* (see Table S1) following the manufacturer’s protocol of the MagAttract HMW Kit (Qiagen). Then, we built Illumina libraries from these DNA extracts and sequenced aiming for ∼20x of coverage in a 2x150 bp NovaSeq6000 lane. For the reference genome, we extracted genomic DNA from the blood of an *Iberolacerta aurelioi* individual (IA003) from Massís de Comapedrosa (Andorra), to sequence a total of two 8M SMRT HiFi PacBio cells (long reads, 34 Gbp) and Illumina short reads aiming for ∼30x coverage (27 Gbp). One Hi-C library was prepared from blood collected from the same individual using the Hi-C High coverage kit (Arima Genomics), following manufacturer’s instructions including custom modifications when necessary. Sample concentration was assessed by Qubit DNA HS Assay kit (Thermo Fisher Scientific) and library preparation was carried out using the ACCEL-NGS® 2S PLUS DNA Library Kit (Swift Bioscience). Library amplification was carried out with the KAPA HiFi DNA polymerase (Roche). Finally, we sequenced the Hi-C in a 2x150 Illumina NovaSeq6000 lane aiming for ∼60x of coverage (52 Gbp). Finally, another *I. aurelioi* specimen from the same locality was sacrificed to extract RNA from five different tissues (liver, eyes, kidneys, heart, and brain) with the HigherPurityTM Tissue Total RNA Purification Kit (Canvax) and quality-checked with Bioanalyzer. RNA extractions were pooled and sequenced with PacBio long reads (Iso-Seq) to obtain HiFi full-length transcriptomes. This sample was sequenced with one eighth of a PacBio HiFi SMRT Cell 8M using the Sequel II sequencing kit (Novogene Co).

### Genome assembly and annotation

To initially explore the genome sequenced data, we generated a k-mer profile with Meryl (Rhie et al., 2020) and we visualized it with GenomeScope2 (Ranallo-Benavidez et al., 2020). Then, we assembled the genome following the VGP assembly pipeline v2.0 (Rhie et al., 2021). PacBio HiFi reads were assembled into contigs using hifiasm (Cheng et al., 2021), producing primary and alternate assemblies. We used purgedups (Guan et al., 2020) to remove haplotypic duplicates from the primary assembly and added them into the alternate. Then, we scaffolded the primary assembly using the Hi-C data with SALSA2 (Ghurye et al., 2017). Manual curation was performed with PretextView (Harry, 2022). We used the ∼30x Illumina data to polish the assembly with Pilon (Walker et al., 2014). Additionally, we identified repeat elements using RepeatModeler v.2.0.3 (Smit & Hubley, 2008) for *de novo* predictions of repeat families. To annotate genome-wide complex repeats, we used RepeatMasker v.4.1.3 (Smit et al., 2013) with default settings to identify known Tetrapoda repeats present in the curated Repbase database (Bao et al., 2015). Then, we ran three iterative rounds of RepeatMasker to annotate the known and unknown elements identified by RepeatModeler. The mitochondrial genome was obtained with MitoFinder (Allio et al., 2020), using the mitochondrial genome of *Podarcis muralis* (Andrade et al., 2019) as a reference.

Quality assessment and general metrics for the final assembly were carried out with gfastats (Formenti et al., 2022). Possible contaminations were evaluated with BlobToolKit (Challis et al., 2020) using the NCBI taxdump database. We also used MitoFinder to confirm that the mitochondrial genome was absent in the assembled nuclear reference genome. Completeness of the genome assembly was assessed with BUSCO v5.3.0. against the sauropsida_odb10 database (n=7,480) (Simão et al., 2015).

We annotated the reference genome of *I. aurelioi* by combining three annotation pipelines: 1) First we used GeMoMa v1.9 (Keilwagen et al., 2016), which is based on the conservation of intron positions in related organisms. As input for GeMoMa, we downloaded the annotated reference genomes of *Podarcis muralis* (NCBI RefSeq Assembly ID: GCF_004329235.1; Andrade et al., 2019), *Podarcis raffonei* (GCF_027172205.1; Gabrielli et al., 2023), *Pantherophis guttatus* (GCF_001185365.1; Ullate-Agote et al., 2014), *Pogona vitticeps* (GCF_900067755.1;Georges et al., 2015), *Lacerta agilis* (GCF_009819535.1) and *Zootoca vivipara* (GCF_011800845.1; Yurchenko et al., 2020). 2) Based on *de novo* sequenced full-length transcriptomes of different tissues. Subreads from Iso-Seq PacBio were first converted into circular consensus sequences (CCS) with the ccs function from pbccs v6.4 (Pacific Biosciences of California, Inc), followed by demultiplexing and primer removal with lima v2.7.1 (Pacific Biosciences of California, Inc). Non-chimeric full-length (NCFL) transcripts were then obtained by removing concatemers with isoseq3 v3.8.3 (Pacific Biosciences of California, Inc) and trimming poly-A tails with the python script tama_flnc_polya_cleanup.py (https://github.com/GenomeRIK/tama/wiki) mapped to each reference genome assembly with the splice-aware mapper pbmm2 v1.10 (Pacific Biosciences of California, Inc). The resulting mapped reads were collapsed into unique isoforms with isoseq3 v3.8.3 with the --do-not-collapse-extra-5exons flag. Collapsed unique isoforms were then used to identify protein-coding regions with GeneMarkS-T (S. Tang et al., 2015). 3) Based on external protein databases following BRAKER2 pipeline (Brůna et al., 2021). We ran BRAKER2 using the ortholog database Sauropsida OrthoDB v10 (Kriventseva et al., 2019) as input to create gene sets with GeneMark-EP+ to train Augustus. Finally, we integrated the different annotation files with TSEBRA (Gabriel et al., 2021), keeping all predictions from GeMoMa and GeneMarkS-T.

### Data processing

Raw Illumina reads for the 12 *Iberolacerta* samples sequenced in this work, and four Lacertidae outgroups (*Zootoca vivipara*: GenBank accession number JAATIO000000000.1; *Podarcis muralis*: SIRZ00000000.1; *Lacerta viridis*: OFHU00000000.1; *Lacerta bilineata*: OFHV00000000.1) were first processed with fastp (Chen et al., 2018) applying a sequence quality filtering of 30, discarding poly-G and poly-X tails, and implementing base correction and adaptor trimming. Filtered Illumina reads were then mapped to the *de novo* assembled reference genome with bwa v.0.7.17-r1188 (H. Li, 2013), PCR duplicates were marked and removed with PicardTools v3.0.0 (https://broadinstitute.github.io/picard/) and alignment files were indexed with SAMtools v.1.10 (Danecek et al., 2021). Before variant calling, *I. aurelioi* reference’s genome sample and outgroups were downsampled to ∼15x coverage to ensure comparability with all other *Iberolacerta* samples. Variant calling was performed independently for each chromosome, excluding sexual chromosome 7, with GATK’s HaplotypeCaller, GenomicsDBImport and GenotypeGVCFs tools (McKenna et al., 2010) applying a minimum base quality score of 30. A strict filtering pipeline was then applied to variant calls to ensure high call reliability. Calls were purged out if FS > 20, QD < 2, MQ < 40, MQRankSum or ReadPosRankSum < -0.5. Indels were normalized and SNPs within ten bp to the indels were excluded. Finally, we discarded genotypes within the inferred repetitive regions and concatenated each filtered autosome into a single vcf with VCFtools (Danecek et al., 2011) which resulted in a final dataset of 80,270,080 SNPs.

### Phylogenomic reconstructions and introgression patterns

First, we explored the genomic relationships among the *Iberolacerta* lineages by means of a Principal Component Analysis (PCA) implemented in Plink v2.00a2.3 (Chang et al., 2015). We filtered the initial dataset by keeping biallelic SNPs present in 90% of the samples and applying a minor allele frequency of 0.05. Then, we assessed the decay of linkage disequilibrium with PopLDdecay (Zhang et al., 2019) and selected one SNP for every 10-kb window to obtain unlinked SNPs (uSNPs), resulting on a final dataset of 125.63k uSNPs. The PCA results were plotted in ggplot2 (Wickham, 2016). Later, to determine not only the phylogenomic relationships within this genus, but among the other Lacertidae outgroups mentioned above (i.e., *Podarcis*, *Zootoca* and *Lacerta*) as well, we performed phylogenomic reconstructions based on genome-wide nuclear SNPs. We first reconstructed a Maximum-Likelihood (ML) phylogeny with nuclear SNPs, in which we filtered the 80.27M SNP dataset with VCFtools excluding SNPs with more than 20% missing data, selected one allele at random at each site, and resulted in a final dataset of 67,673,350 SNPs. Then, we ran IQtree v2 (Nguyen et al., 2015) on 100-kbp non-overlapping windows with a GTR+ASC model (accounting for ascertainment bias), and 100 starting trees. Best trees for each analysis were combined and a consensus tree for all windows was obtained in ASTRAL-III (Zhang et al., 2018).

A Multispecies Coalescent time-calibrated species tree was also inferred with SNAPP (Bryant et al., 2012) implemented on BEAST2 (Bouckaert et al., 2019). We excluded one of the *I. aurelioi* redundant samples and filtered the initial SNP dataset by applying a minor allele filter of 0.05, keeping biallelic SNPs present in at least 90% of the samples, and selecting one SNP every 10 kbp, resulting in a final dataset of 123.8k uSNPs. Following the SNP-based molecular clock of (Stange et al., 2018) we used the “snapp_prep.rb” script (https://github.com/mmatschiner/tutorials) to generate the input file for SNAPP. We constrained the deepest node of the phylogeny with a normally distributed prior centered at 37.55 mya and a standard deviation of 4 according to mean and 95% HPD interval in Garcia-Porta et al., (2019). We then selected 50k uSNPs at random and ran three independent runs of five million generations of SNAPP, sampling every 50 generations. Convergence across runs and stationarity was checked with Tracer v1.7 (Rambaut et al., 2018). Posterior distributions were combined with LogCombiner v.2.6.4, discarding 10% of the posterior trees as burn-in and a maximum clade credibility tree was obtained calculating median heights in TreeAnnotator v2.6.4 (Bouckaert et al., 2019).

To account for possible introgression events among the *Iberolacerta* lineages, we ran ABBA-BABA tests and estimated the *f_4_-ratio* with the function Dtrios from Dsuite v0.5 r53 (Malinsky et al., 2021). First, a dataset with all *Iberolacerta* specimens and outgroups previously used, was filtered with VCFtools to keep only biallelic SNPs below 10% missingness and quality over 30, keeping a total of 53.08M SNPs. Afterwards, Dtrios was ran with the flag ABBA-clustering, to account for the confounding effect of different mutation rates among diverging species (Koppetsch et al., 2023). We only considered significant those trios below the sensitive clustering threshold at α=0.01. Finally, using the inferred topology from the previous analyses, we summarized the *f_4_-ratio* estimates by computing the *f-branch* statistics from Dsuite, which better interprets correlated allele sharing excess across a phylogeny.

### Genetic diversity

For each sample, including the outgroups, we calculated its average genome-wide heterozygosity. For that, we generated non-overlapping 100-kbp sliding windows across the newly assembled reference genome of *I. aurelioi* and took only sites (both variant and invariant) with site quality above 30. Only windows containing more than 60,000 unfiltered sites were considered. Visualization was carried out with ggplot2 (Wickham, 2016).

Afterwards, we explored how different *Iberolacerta* species are affected by Runs of Homozygosity (ROHs). Using the same dataset. ROHs were calculated based on the density of those remaining heterozygous sites by the implemented hidden Markov model (HMM) in bcftools roh function (Narasimhan et al., 2016), setting the allele frequency as 0.4 per default, being thus conservative when identifying ROHs (Dyson et al., 2022). We kept ROHs with a PhredScore above 70 and a minimum length of 100 kbps (Mochales-Riaño et al., 2023). Finally, we divided the amount of ROHs in each high-coverage genome by size, as different-sized ROHs arise from different causes (Ceballos et al., 2018), in the following categories: short (0.1-0.5 Mbp), medium (0.5-1 Mbp) and long (>1 Mbp); and plotted the result with ggplot2, showing the amount of ROHs as a percentage of the total autosomal genome (1.35 Gb).

### Demographic history

We inferred the demographic history of all the species within the *Iberolacerta* genus by implementing the Pairwise Sequential Markovian Coalescent (PSMC) software (Li & Durbin, 2011). Heterozygous positions were obtained from bam files with Samtools v1.9 (Danecek et al., 2021) and data was filtered for low mapping (<30) and base quality (<30). Minimum and maximum depths were set at a third (4x) and double (30x) the average coverage, respectively. Only pseudochromosome scaffolds, excluding sexual chromosome 7 (Z_1_), were considered. A rate of 2.4x10^-9^ substitutions/site/generation (Green et al., 2014) and a generation time of 8 years were used (Mouret et al., 2011). A total of ten bootstraps were calculated and we plotted the results in R, highlighting the diversification of the genus *Podarcis* (4-6.5 mya; Yang et al., 2021) and the Last Glacial Period.

### Macrosynteny analysis

Chromosomal rearrangements were explored with MCscan (H. Tang et al., 2008) among *Iberolacerta aurelioi*, two other lacertid species (*P. muralis*, *L. agilis*), and one amphisbaenian as outgroup: *Rhineura floridana* (Family Rhineuridae; NCBI Accession number: GCF_030035675.1). Protein sequences from each species were extracted using AGAT v0.8.0 (Dainat, 2023) and pairwise aligned using LAST (Kiełbasa et al., 2011) with the JCVI python module (H. Tang et al., 2015). Initial alignments between species were used to identify sexual chromosomes and chromosomes assembled in the reverse complement, which were corrected in some of the assemblies by reverse complementing the fasta sequence with SAMtools faidx v0.7.2.2. Afterwards, GeMoMa was used to generate a draft annotation of the new genomes with reversed sequences, using as reference the original reference genome of each respective species. Then, we re-ran the MCscan pipeline to generate final re-oriented synteny plots.

### Hemoglobin evolution

To identify hemoglobin gene clusters in *I. aurelioi* and other Lacertidae species, we extracted and translated with AGAT v1.0.0 script *agat_sp_extract_sequences.pl* (Dainat, 2023) all annotated coding regions in *I. aurelioi* genome, as well as in *Z. vivipara*, *P. muralis* and *L. agilis* reference genomes, and performed genome-wide protein blasts (Johnson et al., 2008) against α- and β-chains of *P. muralis* hemoglobin sequences (respective accession codes: A0A670JFV3_PODMU and A0A670ICR7_PODMU). We kept hits above a stringent e-value of 10^-8^. Among different isoforms per gene, we chose the one with the best scoring e-value for each of the selected genes. Such translated transcripts were blasted online against the standard blastp database to identify each gene/transcript as α-A, α-D or β hemoglobin subunits, another cytoglobin or a false positive blast result. The chromosomal location of the α- and β-globin gene clusters in *Iberolacerta*, *Lacerta* and *Podarcis* was checked with previous macrosynteny results. Hemoglobin genes were coded from 5’ to 3’ agreeing with *Iberolacerta*’s genome. CDS and protein sequences from hemoglobin genes were aligned in MEGA v11.0.13 (Tamura et al., 2021) with MUSCLE algorithm (Edgar, 2004). We manually inspected for premature stop codons and removed sequences exceeding known hemoglobin lengths (142 and 147 amino acids for α- and β-chains, respectively). We constructed a Maximum Likelihood protein tree with IQtree v1.6.12, and support for nodes was evaluated with 10,000 pseudoreplicates of ultrafast bootstrap procedure, with JTT+G4 chosen as best-fit model according to BIC. Tree results were used to infer orthology relationships between hemoglobin genes within each α- and β gene clusters among the four genera. Expression levels within these two clusters in *Iberolacerta* were explored in Geneious v2023.2.1 in terms of Iso-Seq coverage, log-transformed and normalized independently for each gene cluster.

Finally, we explored selection on the hemoglobin-encoding genes in the mentioned Lacertidae. The CDS alignment of all hemoglobin sequences for the four species was used to test d_N_/d_S_ ratio tests with FEL (Kosakovsky Pond & Frost, 2005) and RELAX (Wertheim et al., 2015) models implemented on Datamonkey 2.0 (Weaver et al., 2018). FEL was tested for the crown groups of α- and β-chain gene clusters, separately, to investigate codons under purifying or diversifying selection at each group. On the other hand, four comparisons were tested with RELAX to check for relaxation or intensification of the selection: 1) α-D *vs* α-A isoforms, 2) *Iberolacerta*’s α-A *vs* other Lacertidae α-A, 3) *Iberolacerta*’s α-D *vs* other Lacertidae α-D and 4) *Iberolacerta*’s β*-*globins *vs* other Lacertidae β-globins.

## Results

### Genome assembly and macrosynteny analysis

We assembled and annotated a high-quality chromosome-level assembly for the Aurelio’s rock lizard (*I. aurelioi*) by combining PacBio HiFi (∼30x), Hi-C (∼60x) and Illumina (∼30x) libraries. Final assembly size is 1.44 Gb, with 13 scaffolds or pseudochromosomes comprising the 98.05% of the assembly (Fig. S1). Scaffold and contig N50 reach, respectively, 172.8 Mbp and 71.4 Mbp. Up to 95.1% of the 7,480 Sauropsida orthologs were found (93.2% as single-copy genes) in the assembly. GC content yielded 43.65% and repeat content 35.09%.

### Phylogenomic reconstructions and introgression events

Within the studied *Iberolacerta*, the genomic PCA splits four species clusters associated to different geographical ranges: Alps (*I. horvathi*), Pyrenees (subgenus *Pyrenesaura*: *I. aranica, I. bonnali,* and *I. aurelioi*), Central System (*I. cyreni cyreni* and *I. c. castiliana*), and northwestern Iberia (*I. martinezricai*, *I. galani*, *I. monticola monticola* and *I. m. astur*) (Fig. 1D). These four groups are supported as well by phylogenomic analyses (Figs. 2A, S2-3), that suggest old origins for most of the studied *Iberolacerta* species. The Alpine species, *I. horvathi*, is the earliest diverging taxon within the genus, separated ∼16.05 mya, followed by the *Pyrenesaura* clade, 13.98 mya (Fig. S3). The split between Central and Northwestern clades, albeit younger, is relatively old as well: 11.25 mya. Indeed, most species-level divergences within *Iberolacerta* occurred during the Miocene. However, *I. martinezricai* and *I. galani* appear to have diverged from the ancestral *I. monticola* more recently, during the Pleistocene (approximately 0.93 and 0.56 million years ago, respectively). Notably, the divergence of *I. galani* occurred around the same time as the split between subspecies within *I. cyreni* (0.50 million years ago), whereas *I. martinezricai* represents an earlier and more distinct lineage within the Pleistocene diversification. Subsequently, *I. monticola* gave rise to its two currently recognized subspecies, which exhibit only shallow genomic divergence. In terms of phylogenetic placement within Lacertidae, *Lacerta* is more closely related to *Iberolacerta* than any of the other genera analyzed, with *Zootoca* being more distantly related than *Podarcis* (Fig. 2A).

**Fig. 2.**
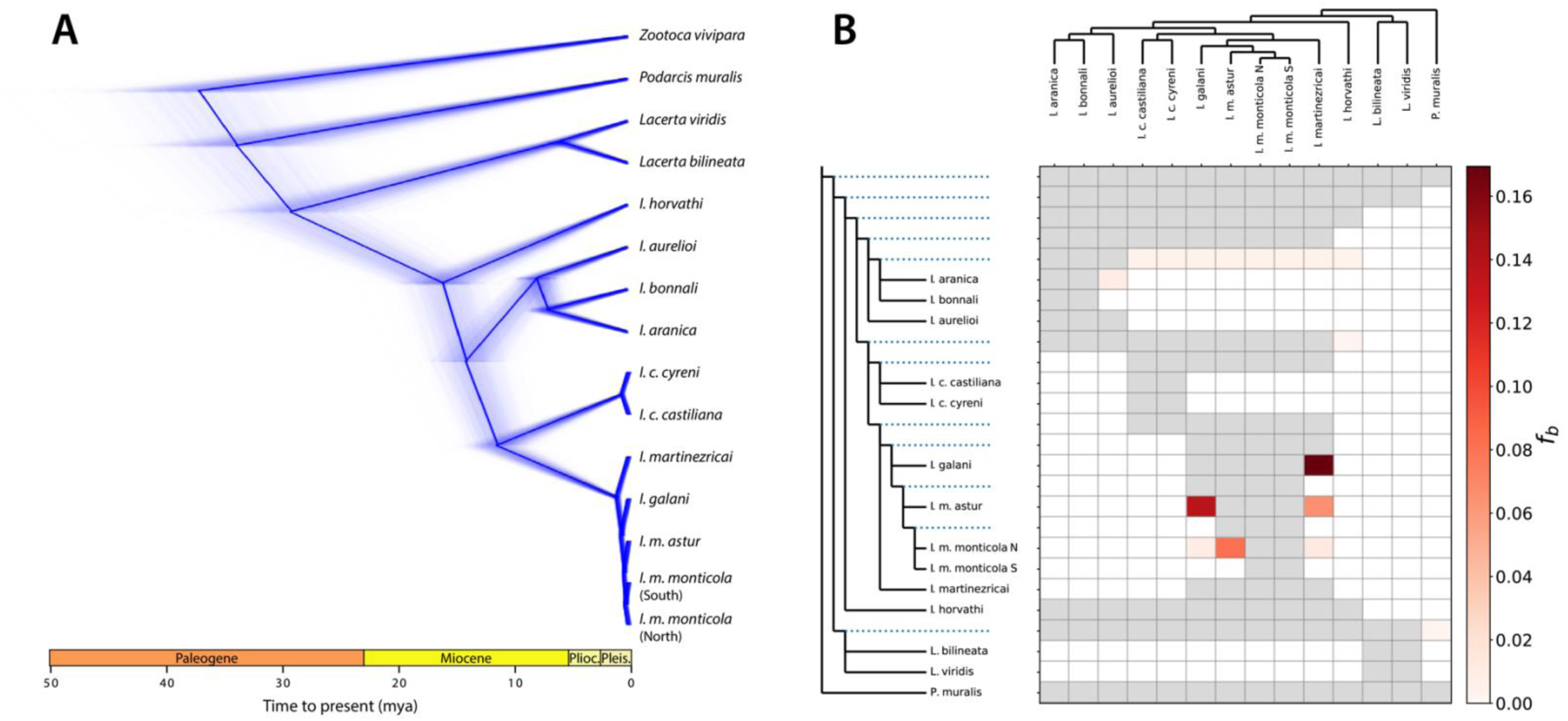
Evolutionary relationships of the genus *Iberolacerta*. A) Genomic phylogeny of the genus *Iberolacerta*, including other Lacertidae genera as outgroups, performed with SNAPP. *Iberolacerta* is closer to *Lacerta* than to *Podarcis* or *Zootoca*. Within *Iberolacerta*, the divergences among the four main clades occurred during the Miocene. The alpine species, *I. horvathi*, is the first branching out, followed by the subgenus *Pyrenesaura*, comprised of I*. aranica, I. bonnali,* and I*. aurelioi*. Within the sister Central System (*I. cyreni* sspp.) and NW Iberian clades (*I. martinezricai*, *I. galani*, and *I. monticola* sspp.) we found subspecific diversity of Pleistocene origin, as well as the divergence of *I. martinezricai* and *I. galani*. Node ages with 95%HPD are displayed in Fig. S3. All nodes have posterior probabilities equal to 1. B) Excess of allele-sharing (*f-branch* statistics) to measure introgression. Within the NW Iberian clade of *Iberolacerta*, introgression is widespread and extensive, with up to >16% of the diversity of *I. galani* and *I. martinezricai* being shared.

Concerning introgression events, a high excess of allele sharing was detected among several pairs of the NW Iberian set of *Iberolacerta* lineages (Fig. 2B): between *I. galani* and *I. martinezricai* (16.94% of shared diversity); and between *I. monticola astur* and other related taxa: *I. galani* (13.76%), *I. monticola monticola* from the Cantabrian Mts. (8.14%), and *I. martinezricai* (6.53%).

### Demography, heterozygosity and ROHs

Demographic inference results show deep bottlenecks in Alpine and, especially, Pyrenean *Iberolacerta*, during the Pliocene, matching the major diversification of the genus *Podarcis* (Yang et al., 2021) in the Iberian Peninsula and the Mediterranean region (Fig. 3A). On the other hand, NW Iberian and Central System species underwent only modest decline after *Podarcis* diversification, but moderate population expansions occurred during the Pleistocene, especially for *I. monticola*. However, all these Iberian populations contracted around the Late Pleistocene (Fig. 3A). Regarding genome-wide heterozygosity and ROH burden (Fig. 3BC), *Iberolacerta* species follow two different patterns. On the one hand, the Alpine and Pyrenean species yielded low heterozygosity values and large percentages of their genome covered by ROHs, including long ones. Both shorter and longer ROHs point towards ancient bottlenecks and current inbreeding (Ceballos et al., 2018), matching their prolonged histories with exiguous population sizes. Second, Central and NW Iberian species show similar or even higher mean heterozygosity values than other lacertids, and their ROH burden is composed mainly by short ROHs, in agreement with Pleistocene population contractions, especially for *I. cyreni* and *I. m. monticola*, whose proportion of short ROHs is comparable to the Alpine and Pyrenean counterparts.

**Fig. 3.**
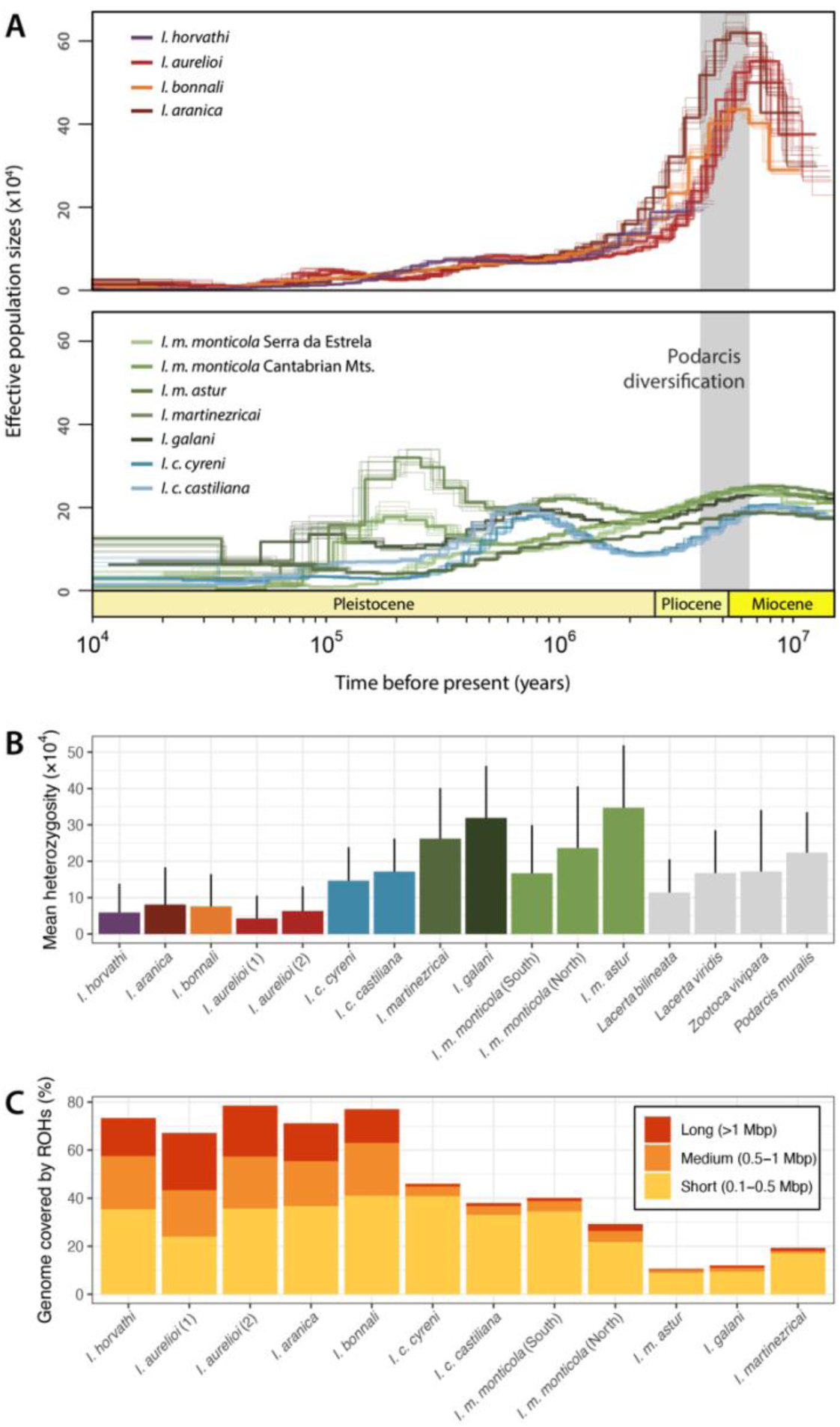
Demographic oscillations, heterozygosity estimates and ROH burden. A) Alpine and Pyrenean species (top) suffered severe bottlenecks around the Pliocene, coinciding with the diversification of *Podarcis* (highlighted in light grey), whereas NW Iberian and Central System species (bottom) underwent only modest population declines around the Pliocene and populations expansions during the Pleistocene, followed by subsequent contractions. B) Heterozygosity is shown as genome-wide means + standard deviations per individual and ROH burden as the percentage of the genome covered by runs of homozygosity, classified by their length (C). Alpine and Pyrenean species showcase lower heterozygosity estimates and greater ROH burden.

### Chromosomal rearrangements

Although the genome size of *I. aurelioi* is similar to that of other lacertids (Andrade et al., 2019; Gomez-Garrido et al., 2023), its haploid number is significantly reduced compared to the ancestral Lacertidae karyotype, depicted by *Lacerta* and *Podarcis* genomes (Fig. 4). Six out of 13 chromosomes in *Iberolacerta* are the result of chromosome fusions, highlighting the one between the ancestral Z chromosome and the only microchromosome typically found in lacertids but absent in *Iberolacerta*. This fusion gave rise to the Z_1_ chromosome (chr. number 7; Fig. 4).

**Fig. 4.**
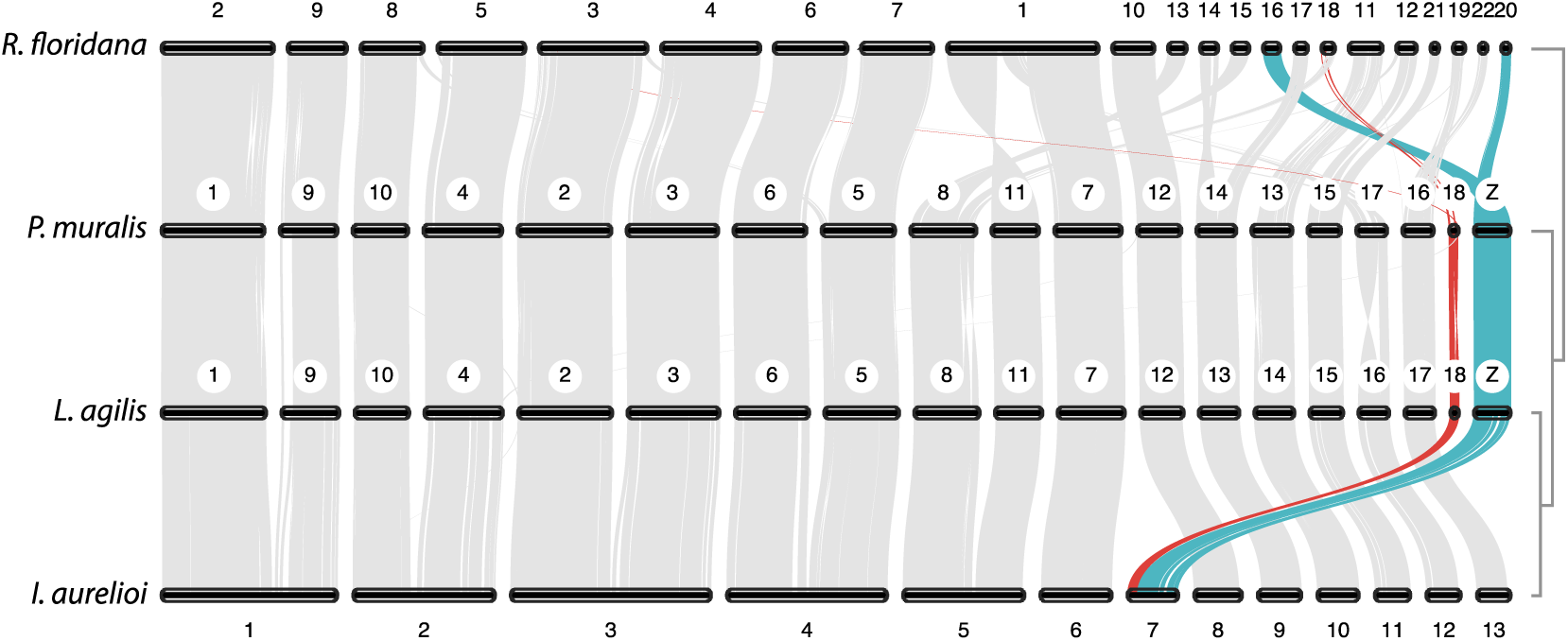
Macrosynteny analysis showing chromosomal rearrangements in *Iberolacerta* and other Lacertidae. *Rhineura floridana*, an amphisbaenian, sister clade to Lacertidae, was included as outgroup. Major chromosomal reorganization has occurred between Lacertids and amphisbaenians. Within Lacertids, karyology seems better conserved, with 17 autosomal macrochromosomes and only one microchromosome, as depicted by *Podarcis muralis* and *Lacerta agilis*. *Iberolacerta aurelioi* exhibits a much lower haploid number (n=13), due to six chromosomal fusions (chrom. 1-5 and 7; and an extra fusion for females). One of those fusions affects the ancestral Z chromosome (turquoise) and the only microchromosome (red), giving rise to the Z_1_ chromosome. Chromosome Z_2_ has not been identified.

### Hemoglobin evolution

The α and β Hemoglobin (Hb) subunits appear to be well conserved from a macrosyntenic point of view: chromosomal locations of both *Iberolacerta’s* Hb clusters match Hb loci in *Podarcis* and *Lacerta*, despite the chromosomal rearrangements observed in *Iberolacerta*. Regarding microsynteny, the β-globin cluster displays the same architecture as the closely-related genus *Lacerta*, while the α-cluster is well conserved throughout the phylogeny, as depicted by the genera *Podarcis* and *Zootoca*, and in spite of a more derived configuration in *Lacerta* (Fig. 5A). All three β-chain genes and four out of five α-chains are expressed in *Iberolacerta*. Hemoglobin α-subunit comprises the isoforms A and D, being the latter the major isoform in squamates, although its affinity for oxygen is lower than isoform A (Hoffmann et al., 2018). In *Iberolacerta aurelioi*, Hb α-A, of enhanced oxygen affinity, shows a slightly higher expression (Fig. 5B). Concerning sequences, the two clades of α-chains and the β-chains are reciprocally monophyletic. *Iberolacerta* Hb proteins do not show longer branches than other Lacertidae species (Fig. 6A). Fixed Effects Likelihood (FEL) models found no codons under positive diversifying selection for Hbα, and only one site for Hbβ at p≤0.05. However, purifying selection was found in 31 and 32 codons at each Hb group at p≤0.05, respectively. Such selection appears to have been significantly intensified in all α-D isoforms, in regard to α-A (K=4.62; p-value=0.006; Fig. 6B). However, *Iberolacerta*’s α-A genes have suffered the greatest intensification on their selection compared to the Hb-encoding genes in *Podarcis*, *Lacerta* and *Zootoca* (K=21.25; p-value=0.015; Fig. 6B). On the other hand, neither *Iberolacerta*’s α-D nor β subunits have undergone significant intensification nor relaxation compared to other Lacertidae.

**Fig. 5.**
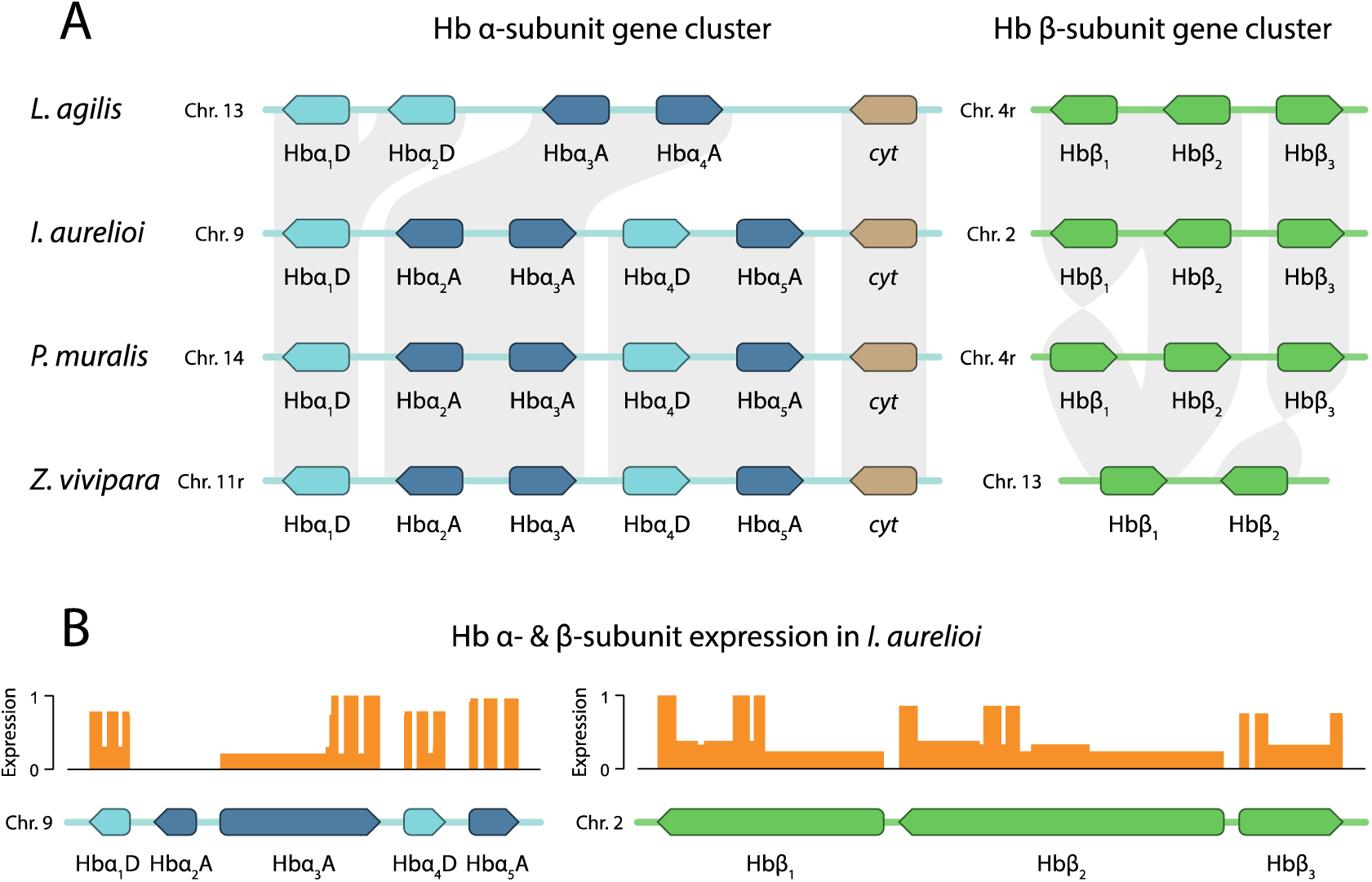
Microsynteny and expression levels of hemoglobin (Hb) gene clusters. A) Microsynteny comparisons of the gene clusters encoding α and β subunits of Hb in *Iberolacerta* and other Lacertidae. Cyt stands for a cytoglobin (brown) not related to hemoglobin. Chromosomes annotated with “r” have been depicted with its corresponding reverse complement. Microsynteny of Hbα (blue) cluster in *Iberolacerta* is equal to *Podarcis* and *Zootoca*, whereas Hbβ (green) is equal to *Lacerta*. B) Expression levels of each gene within each Hb subunit cluster, in terms of relative Iso-Seq coverage. Isoform A (dark blue) of subunit α, of enhanced oxygen affinity, is slightly more expressed in *I. aurelioi* than isoform D (light blue), which is, in contrast, the major isoform in Squamates.

**Fig. 6.**
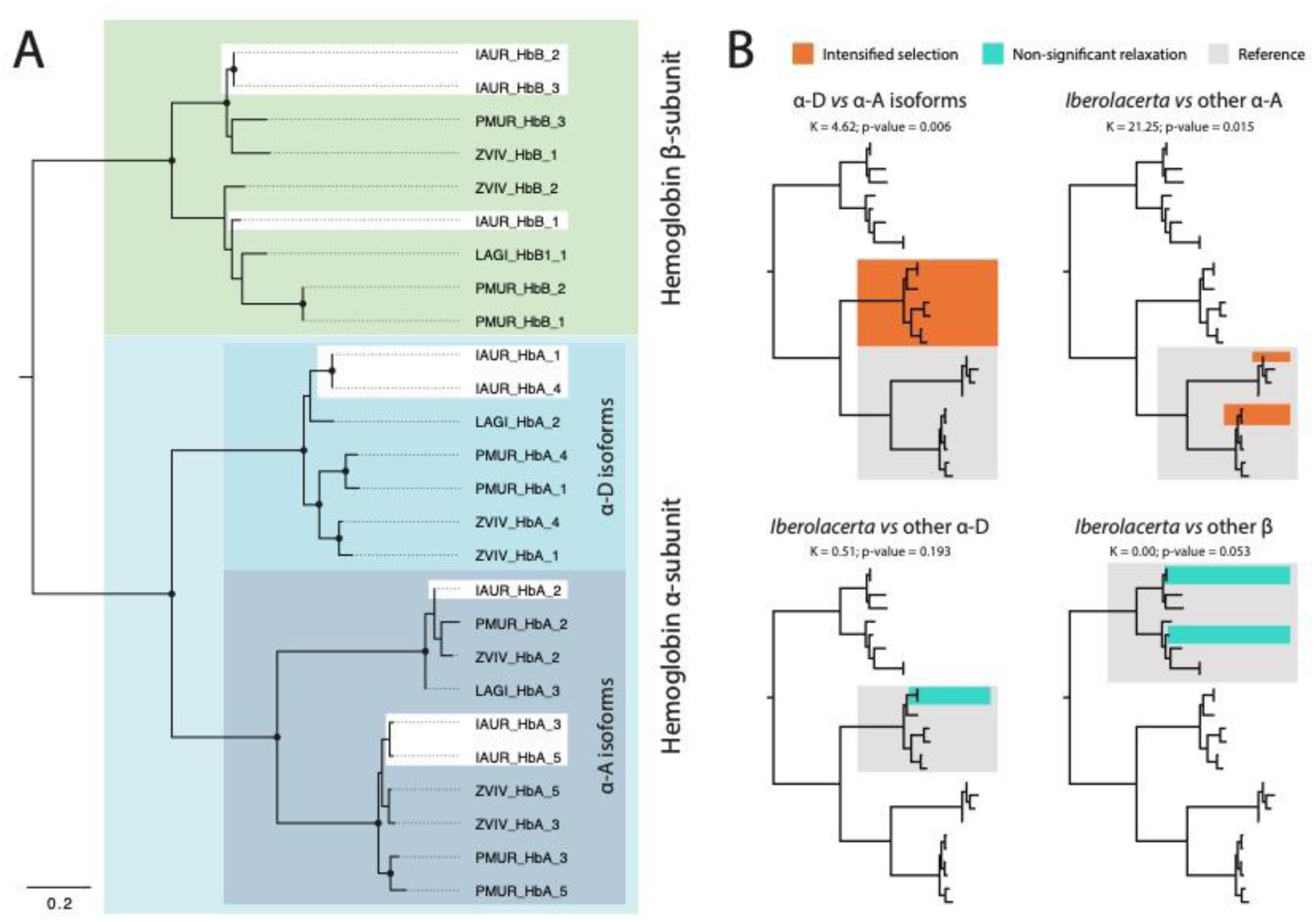
Selection analyses on hemoglobin genes. A) Hemoglobin protein tree, including α and β subunits found in *Iberolacerta aurelioi* (IAUR), *Lacerta agilis* (LAGI), *Podarcis muralis* (PMUR) and *Zootoca vivipara* (ZVIV). *Iberolacerta* hemoglobins show short branches. B) RELAX test results on hemoglobin CDS. Purifying selection has been intensified in D isoforms *vs*. A isoforms of Hbα, in line with D being the major isoform in Squamata. However, purifying selection has been more intensified in *Iberolacerta* Hbα-A isoforms compared to A isoforms of other lacertids, in line with a higher expression of this putatively minor isoform of enhanced oxygen affinity.

## Discussion

Our phylogenomic results support the segregation of the genus *Iberolacerta* in four old clades, from East to West and from older to younger: Alps, Pyrenees, Central System and NW Iberia, in agreement with previous nuclear and mitochondrial DNA studies of the genus (Arribas et al., 2006; Carranza et al., 2004; Garcia-Porta et al., 2019). However, some mito-nuclear discordance is found, as studies based on mitochondrial genes had suggested different relationships among the four clades, albeit with low support, possibly due to rapid divergences among them (Carranza et al., 2004). Our time-calibrated species tree confirms an ancient diversification of this genus in the Early Miocene (16.7 mya, Fig. 2A, S3), backing up genomic studies on Lacertidae (14.6 mya in Garcia-Porta et al., 2019) but in contrast to previous multi-locus works, which suggested a much younger origin (7.7 mya in Carranza et al., 2004).

Despite the old origin of the clades, and even old species divergences within clades (e.g., *I. aurelioi* split from other *Pyrenesaura* during the Miocene, 7.82 [6.16-8.42] mya, Fig. S3), those long “bald” branches either have not undergone further diversification processes until the recent Pleistocene, in line with a relict nature of *Iberolacerta* rock lizards (Carranza et al., 2004), or evidence the extinction of lost ancient lineages, probably throughout the glacial cycles. During the Pleistocene, population expansions, followed by contractions, are reported for NW Iberia and Central System taxa (Fig. 3A), in agreement with shallow genomic differentiation occurring in the mid and late Pleistocene (Fig. 2A), and the occurrence of introgression events among the former clade species (Fig. 2B). Particularly interesting is the low differentiation between the disjoint Portuguese Central System and the Cantabrian populations of *I. monticola*, despite the ∼200 km separating Serra da Estrela (Central Portugal) from the nearest population of *I. m. monticola* in Galicia (NW Spain). This suggests that Galician populations, especially the now-relict demes by the coast and inland Galicia (Orense, Serra da Queixa), might have extended southwards up to Serra da Estrela through a lowland corridor in North Portugal during glacial maxima. Indeed, part of this corridor still lies within the potential range of the species (Araujo et al., 2011), and microsatellite data has revealed that fragmentation of lowland *I. m. monticola* populations is somewhat related to recent climatic and human-induced loss of Atlantic forests (Remón et al., 2013).

Regarding the NW Iberia clade, the species *I. martinezricai* first, and *I. galani* secondly, branched out during the Pleistocene, the latter at an age that is more similar to the subspecific diversification of *I. cyreni* than to other speciation ages within the genus, and both of them notably more recent than speciation events of other Mediterranean lacertids (e.g. Yang et al., 2021). Despite their recent divergence, cytogenetic evidence supports the long-term isolation of *I. martinezricai*, which shows a slightly different karyotype from both *I. monticola* and *I. galani*. In particular, *I. martinezricai* exhibits a nucleolar organizer region (NOR) located on a medium-sized chromosome pair (MS-NOR), in contrast to the large chromosome pair NOR (L-NOR) found in the latter two species (Arribas & Odierna, 2004). This MS-NOR configuration is otherwise only present in *I. cyreni*, whereas all other *Iberolacerta* taxa share the L-NOR condition, underscoring the distinct evolutionary trajectory of *I. martinezricai*. Given its unique cytogenetic signature, high-elevation specialization, and Endangered status, *I. martinezricai* represents a priority for conservation and further genomic research.

Interestingly, previous multilocus phylogenies, largely based on mitochondrial genes, had suggested a slightly different topology and divergence times. In particular, *I. galani* and *I. martinezricai* were recovered as sister taxa (albeit with low support), diverging around 2 mya and separating from *I. monticola* approximately 2.5 mya (Arribas et al., 2006), in contrast to the more recent genomic estimates of 0.93 and 0.56 mya, respectively (Fig. 2A, S3). Likewise, the divergence of *I. monticola astur* had been previously estimated at 1.8 mya (Arribas et al., 2014; Remón et al., 2013), compared to the much younger estimate of 0.32 mya from our genomic data (Fig. S3). These deeper mitochondrial divergences may reflect long periods of geographic isolation followed by secondary contact (Hogner et al., 2012), particularly if there is strong male-biased dispersal, as documented in *I. cyreni* (Aragón et al., 2001).

In addition, an alternative –and common– source of mito-nuclear discordance is the adaptive introgression of mitochondrial DNA (Toews & Brelsford, 2012), already hypothesized for *I. m. astur* (Arribas et al., 2014). As our results support, an older divergence of *I. galani* (and its uncertain position in the phylogeny) and *I. m. astur* reported by multilocus phylogenies, compared to genomic data, may be explained by the extensive introgressive hybridization with *I. martinezricai* and *I. galani*, respectively, involving not only the 16.9% and 13.7% of their genomic diversity but also their mitochondrial genomes (Fig. 2B). *Iberolacerta martinezricai* and *I. galani* were described as distinct species, apart from morphological and karyological evidence, based on their putatively deep divergence times (Arribas et al., 2006; Arribas & Carranza, 2004; Arribas & Odierna, 2004). Consequently, further research should address whether this diversification within the NW Iberian *Iberolacerta* clade is better interpreted at the subspecific level, especially for *I. galani*.

In contrast to the Pleistocene expansions and diversification of NW Iberian and Central System rock lizards, Pyrenean and Alpine species showed an opposed demographic trend. Despite large populations during the Miocene, they were affected by severe bottlenecks during the Pliocene (Fig. 3A), coinciding with the diversification of the genus *Podarcis* (Yang et al., 2021). This supports the competitive exclusion hypothesis, in which *Podarcis* relegated *Iberolacerta* rock lizards, or at least some species, to exclusively inhabit high-altitude environments (Carranza et al., 2004). These minimal population sizes appear to have been maintained throughout the Pleistocene glacial oscillations up to the present, likely by relying on nearby mountain refugia during glacial maxima (Crochet et al., 2004; Mouret et al., 2011). This demographic stability under constrained conditions has resulted in low heterozygosity values and a high burden of runs of homozygosity (ROH) (Fig. 3B). Quantifying this loss of genetic diversity is critically important for conservation, as it can limit the adaptive potential of populations and jeopardize their long-term viability (Brüniche-Olsen et al., 2018). This concern is particularly acute for the Endangered *I. aurelioi*, which exhibits a notable ROH burden. In contrast, the also Endangered *I. martinezricai* shows little genomic evidence of inbreeding, despite being considered one of the rarest European reptiles, characterized by low wild population densities and a fragmented distribution (Lizana-Ciudad et al., 2021). These contrasting patterns highlight the need for future research to focus on population-level dynamics in these threatened taxa.

Low effective population sizes create the perfect scenario for genetic drift to operate, easily driving to the fixation of chromosomal rearrangements once they appear (Bush et al., 1977; Lande, 1979). In this work we confirm, as previously hypothesized, that the largely reduced diploid numbers found in *Pyrenesaura* species are due to centric Robertsonian fusions between acrocentric chromosomes, giving rise to biarmed metacentric chromosomes, with highly conserved synteny within them (Fig. 4) and without involving insertional translocations (Naveira et al., 2023; Odierna et al., 1996; Olmo et al., 2004). Although other *Iberolacerta* species, such as *I. monticola*, exhibit karyotypes more similar to those of other Lacertidae, the loss of the microchromosome pair was first described by Odierna et al. (1996), and later recognized as a synapomorphy and diagnostic trait of the genus by Arnold et al. (2007) (see also Naveira et al., 2023).Here, we confirm its fusion to the ancestral lacertid Z chromosome to form the new Z_1_ chromosome in *I. aurelioi* (chr. 7; Fig. 4). Although fusion between autosomes and sex chromosomes is a common way to produce neo-sex chromosomes (Kitano et al., 2009; McAllister, 2003), female heterogamety with multiple sex chromosomes (i.e., Z_1_Z_2_W) is rare amongst lizards (3% of studied non-snake Squamata species; Mezzasalma et al., 2021). According to Naveira et al. (2023), this multiple sex chromosome system arose, in the common ancestor of *I. aurelioi* and *I. bonnali*, through the fusion of the ancestral W and one of the autosomes belonging to either pair 15 or 16, resulting in the new W, whereas the homologous chromosome in that pair became Z_2_. Unfortunately, the reference genome of *I. aurelioi* here presented belongs to a male specimen and, therefore, identifying Z_2_ has not been possible. Overall, a total of seven chromosomal fusions set up *I. aurelioi* genome architecture. However, an even lower diploid number has been reported in the Pyrenean *I. bonnali* (2n = 22-24 in males, 23 in females), with one or two additional fusions depending on the population, as intraspecific variation in their karyotypes have been reported (Olmo et al., 2004). While *I. aranica* and *I. aurelioi* share the diploid number (2n = 26), the former lacks the Z_1_Z_2_W system, which was interpreted as a synapomorphy between *I. bonnali* and *I. aurelioi* (Naveira et al., 2023). However, these taxa are not recovered as sister species in this study, likely pointing to convergence in the emergence of the multiple sex chromosome system. Furthermore, these major chromosomal rearrangements among *Pyrenesaura* species may have driven reproductive isolation (e.g. Talavera et al. (2024)), and thus a lack of introgression (Fig. 2B), despite their close geographic and ancient divergence times –unlike the pattern observed in NW Iberian species. Although fusions being the most common chromosomal rearrangement in animals (King, 1995), its prevalence in the genus *Iberolacerta*, particularly in the subgenus *Pyrenesaura*, is remarkable.

Chromosomal fusions physically link genes that were preciously located on separated chromosomes and can lead to reduce recombination rates within the newly formed chromosome arms (Dumas & Britton-Davidian, 2002). Since local adaptation often benefits from reduced recombination between co-adapted loci (Lenormand & Otto, 2000), such fusions are thought to promote adaptation to divergent environments. This mechanism has been documented across multiple taxa, including insects (Bidau et al., 2012), fish (Liu et al., 2022; Wellband et al., 2019) and mammals (Li et al., 2023; Nevo, 2013). Among squamates, there is a general trend towards the loss of microchromosomes through their fusion with macrochromosomes, a process that contributes to the reduction of chromosome number and structural reorganization of the genome (Srikulnath et al., 2021). This trend has been enhanced in Lacertidae, with most species bearing just one pair of microchromosomes (Naveira et al., 2023). Apart from *Iberolacerta*, another cold-adapted lacertid genus has independently lost the microchromosome pair: *Zootoca*, the land reptile with the highest latitudinal distribution (Garcia-Porta et al., 2019; Naveira et al., 2023). Considering the convergence example between the cold-adapted *Iberolacerta* and *Zootoca*, it is conceivable that chromosome fusions might have mediated in adaptation to high altitude environments in *Iberolacerta*, particularly in *Pyrenesaura*, the clade exhibiting the lowest diploid numbers and the highest minimal elevations within the genus. Some of these adaptations might be related to highly effective thermoregulation strategies (Aguado & Braña, 2014; Ortega et al., 2016c, 2016a) and egg retention, as Pyrenean *Iberolacerta* are characterized by an advanced embryonic development at oviposition, which suggests a stage close to viviparity (Arribas & Galán, 2005).

However, chromosome fusion does not only promote local adaptation by linking alleles and reducing recombination. Major gene expression changes have been reported as well owing to Robertsonian fusions through chromatin remodeling, impacting long-distance interactions and chromatin accessibility (Li et al., 2023). Indeed, our results shed light on the adaptation to hypoxic environments in *Pyrenesaura* through gene expression changes, in which a formerly minor Hbα isoform of enhanced oxygen affinity becomes the most expressed (Fig. 5B). A rapid intensification on the purifying selection or constraint on the genes encoding this isoform in *Iberolacerta* (Fig. 6B) backs up its new key role, although, interestingly, this gene cluster is located in one of the few chromosomes not affected by fusions (chr. 9; Fig. 4). Albeit counterintuitively, the development of new traits, such as perhaps hypoxia tolerance in *Iberolacerta*, can be related to increased evolutionary constraints on certain preexisting biological pathways, which become crucial once the new traits are established (Kowalczyk et al., 2020). Overall, this expression shift while conserving the ancestral paralogs and their sequences is consistent with previous findings in Lacertidae, in which adaptation to new environments seems to be characterized by a high degree of genomic reuse (Wollenberg Valero et al., 2022).

Regarding the causes behind the alpine confinement of the genus, both the hyperoxia-as-constraint and the *Podarcis* competitive exclusion hypotheses seem to be backed up by our results in Pyrenean *Iberolacerta*. *Pyrenesaura* species and possibly *I. horvathi* bottlenecks match both the Early Pliocene —a cooling period which should have favored a cold-adapted genus—, and the diversification of *Podarcis* in the Western Mediterranean, although evidence supporting the competitive exclusion by *Podarcis* has only been reported in *I. horvathi* (Žagar et al., 2015). Following mountain confinement, adaptation to hypoxia and other cold-related adaptations likely arose, precluding Pyrenean species to expand their ranges into lowlands during glacial maxima (Fig. 3A). In contrast, both NW Iberian and Central System clades seem to have undergone downslope Pleistocene population expansions, possibly favored by less strict adaptations to hypoxia. This is not surprising, as high-altitude adaptations must occur independently in each clade, since colonization between mountain ranges from a putatively high-altitude adapted ancestor would be unlikely. Indeed, *I. cyreni* has shown plastic adaptive potential to moderate hyperoxia (Megía-Palma et al., 2020), while *I. monticola* elevation range is still wide in the present (Remón et al., 2013). In addition, for these two clades, evidence against the competitive exclusion hypothesis by *Podarcis* has been reported (Galán, 2019; Monasterio et al., 2010), which could be explained, at least in part, by size differences between *Iberolacerta* species groups (e.g., Losos, 1996; Persson, 1985; Schoener, 1983). Whereas NW and Central Iberian *Iberolacerta* species are larger (∼9 cm) than the average *Podarcis muralis* (7.5 cm) (Speybroeck et al., 2016), Pyrenean and Alpine *Iberolacerta* are notably smaller (body length of 6-6.5 cm), which may already represent an adaptation to high altitudes as reported in other lizards (Liang et al., 2023). In absence of *Podarcis* competence, other causes might explain the distributions of NW and Central Iberian species, such as habitat fragmentation (Remón et al., 2013), thermal constraints on embryonic development (Monasterio et al., 2011), or lower refuge availability and thermal quality of lower-elevation habitats (Monasterio et al., 2009).

The genus *Iberolacerta* is probably one of the most studied lizard genera in Europe, attracting remarkable interest from diverse disciplines, such as sex chromosome evolution, thermal ecology, biogeography, or high-altitude adaptation among others. This work aims to advance in some of these research lines by supporting previous hypotheses on the evolution of the genus, like the competitive exclusion triggered by *Podarcis* or the hyperoxia-as-constraint hypotheses, at least in some of its species. The assembly of a high-quality reference genome entails an inflection point for some of those research lines, as well as insights into phylogenomics and ROH burden which will be valuable to systematics and conservation purposes for this threatened genus of rock lizards.

### Data and Material Availability

The genome assembly for *I. aurelioi* and WGS data used in this study will be released upon publication of this manuscript within the GenBank BioProject PRJNA1247761.

### Sampling permits

Sampling of *Iberolacerta aurelioi* for the reference genome sequencing and annotation was authorized by the Departament de Medi Ambient i Sostenibilitat from the Andorran Government with identification code 275628S. Samples used for whole genome resequencing have been used in previous phylogenetic studies on *Iberolacerta* (Mayer & Arribas 2003; Carranza et al. 2004; Arribas & Carranza 2004; Arribas et al. 2006; Arribas et al. 2014) and were authorized by the competent authorities at that time - Departament de Medi Ambient, Generalitat de Catalunya; Consejería de Medio Ambiente, Junta de Castilla y León, Ministeri d’Agricultura i Medi Ambient, Govern d’Andorra; Parque Natural da Serra da Estrela, Portugal; Consellería de Medio Ambiente, Xunta de Galicia; Consejería de Medio Ambiente, Principado de Asturias; and Consejería de Medio Ambiente, Comunidad de Madrid.

## Acknowledgements

We are very grateful to Jordi Solà de la Torre, Cap d’unitat de Fauna, Departament de Medi ambient i Sostenibilitat, Govern d’Andorra for the permit to sample *I. aurelioi,* Eva López, Cap de Medi Ambient, Agricultura, Sostenibilitat i del Parc Natural del Comapedrosa and Jordi Nicolau from Biocom-Also, we must thank Jo Wood and Klara Eleftheriadi for their help during the assembly and curation processes, and Oriol Lapiedra for his support. This study was funded by the Institut d’Estudis Catalans under the Catalan Initiative for the Earth Biogenome Project (Second call for genome sequencing of eukaryotic species from the Catalan territory), by project PID2021-128901NB-I00 funded by MCIN/AEI/10.13039/501100011033 and by ERDF, a way of making Europe, and by grant 2021 SGR 00751 from the Departament de Recerca i Universitats de la Generalitat de Catalunya to SC.AT is supported by “la Caixa” doctoral fellowship program (LCF/BQ/DR20/11790007), BB-C is supported by FPU grant from Ministerio de Ciencia, Innovación y Universidades, Spain (FPU18/04742), GM-R is supported by an FPI grant from the Ministerio de Ciencia, Innovación y Universidades, Spain (PRE2019-088729). ME is supported by an FPI grant from the Ministerio de Ciencia, Innovación y Universidades, Spain PRE2022-101473. HT-C is supported by a “Juan de la Cierva - Formación” postdoctoral fellowship (FJC2021-046832-I) funded by MCIN/AEI/10.13039/501100011033 and by the European Union NextGenerationEU/PRTR, and by a Humboldt Postdoctoral Fellowship from the Alexander von Humboldt Foundation.

