## Supplementary Tables and Figures for "Lizards on a sky archipelago: Genomic approaches to the evolution of the mountain genus *Iberolacerta*"

**Table S1. Samples sequenced in this work with Illumina short reads, the taxon to which they belong, locality and final coverage.** The first specimen was used for reference genome assembly.

| Sample | Species | Location | Coverage |
|---|---|---|---:|
| 003IA | *I. aurelioi* | La Massana (Andorra) | 33.9 |
| SPM004615 | *I. aurelioi* | Pica d'Estats, Pyrenees (Spain) | 15.03 |
| SPM003824 | *I. bonnali* | Bigorre, France | 12.96 |
| 004IAR | *I. aranica* | Val d'Aran, Pyrenees (Spain) | 14.67 |
| LMCYN1S | *I. c. cyreni* | S. de Guadarrama, Central System (Spain) | 14.31 |
| SPM004977 | *I. c. castiliana* | Pico Zapatero, Central System (Spain) | 13.96 |
| E31073 | *I. martinezricai* | Peña de Francia, Central System (Spain) | 12.41 |
| SPM004844 | *I. galani* | Sanabria, Montes de León (Spain) | 13.63 |
| LMO-SE2 | *I. m. monticola* | Serra da Estrela, Central System (Portugal) | 13.69 |
| SPM004603 | *I. m. monticola* | Peña Ubiña, Cantabrian Mts. (Spain) | 14.61 |
| SPM004607 | *I. m. astur* | Catoute Massif, Cantabrian Mts. (Spain) | 15.35 |
| DT80-2 | *I. horvathi* | Carnian Alps, Carinthia (Austria) | 13.32 |

**Table S2. Runs of homozygosity (ROH) burden and genome-wide heterozygosity values per sample.** ROHs are shown as the percentage of the callable genome covered by them, sorted out by length: long (>1Mb), medium (>0.5 Mb & 1 Mb) and short (>0.1 Mb & <0.5 Mb). Heterozygosity values are shown as mean±SD heterozygous sites for every 10 kbp. Metadata for sample codes is found in Table S1.

| **Sample** | **long ROHs** | **medium ROHs** | **short ROHs** | **Heterozygosity** |
|---|---:|---:|---:|---:|
| DT80-2 | 15.92 | 22.15 | 35.27 | 5.91±7.92 |
| 003IA | 23.83 | 19.32 | 23.92 | 4.29±6.27 |
| SPM004615 | 21.29 | 21.62 | 35.62 | 6.30±6.79 |
| SPM003824 | 14.00 | 22.00 | 41.00 | 7.56±8.91 |
| 004IAR | 15.81 | 18.72 | 36.63 | 8.10±10.21 |
| LMCYN1S | 1.16 | 4.03 | 40.81 | 14.66±9.20 |
| SPM004977 | 1.34 | 3.61 | 33.01 | 17.18±9.04 |
| E31073 | 1.40 | 0.90 | 17.00 | 26,22±13.88 |
| SPM004844 | 1.043 | 1.51 | 9.46 | 31.90±14.29 |
| LMO-SE2 | 1.35 | 4.29 | 34.46 | 16.70±13.24 |
| SPM004607 | 0.50 | 1.19 | 8.87 | 34.72±17.16 |
| SPM004603 | 2.90 | 4.55 | 21.75 | 23.64±16.93 |
| *L. bilineata* | - | - | - | 11.41±9.13 |
| *L. viridis* | - | - | - | 16.76±11.78 |
| *Z. vivipara* | - | - | - | 17.21±16.88 |
| *P. muralis* | - | - | - | 22.39±11.12 |

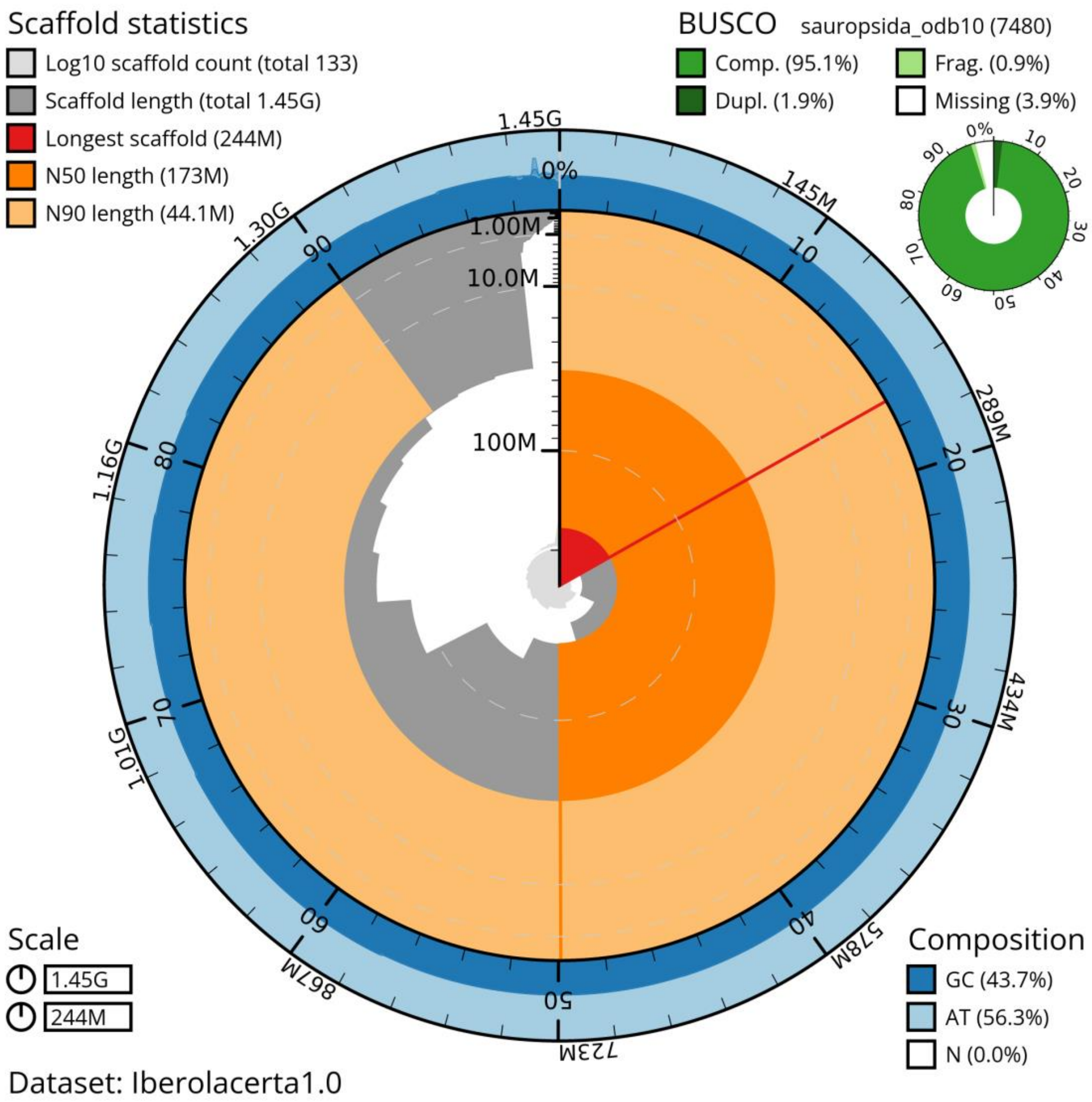


**Fig. S1. Snail plot of *Iberolacerta aurelioi* reference genome assembly presented in this study, summarizing some contiguity and completeness metrics.** M=Mega bases; G=Giga bases. The scaffolds contained in the assembly are shown ordered by length in the inner circle of the snail plot. The sequences within the N50 (173 Mb) and N90 (44.1 Mb) length marks are painted in dark or light orange, respectively. In the outer circle, the GC content of the sequences in represented in blue, whereas AT content in pale blue. Top left corner shows contiguity statistics like the longest scaffold (244 Mb) or Scaffold length (total 1.6 Gb) which indicates the assembly size. Top right corner represents the assembly genetic completeness using BUSCO scores with odb10 Sauropsida database with 7,480 genes, with 95.1% of the database genes found as Single-Copy Complete genes. Bottom left has a legend with the length in bases represented in the snail plot, the perimeter (1.45 Gb) and the radius (244 Mb, i.e., chromosome 1 length).

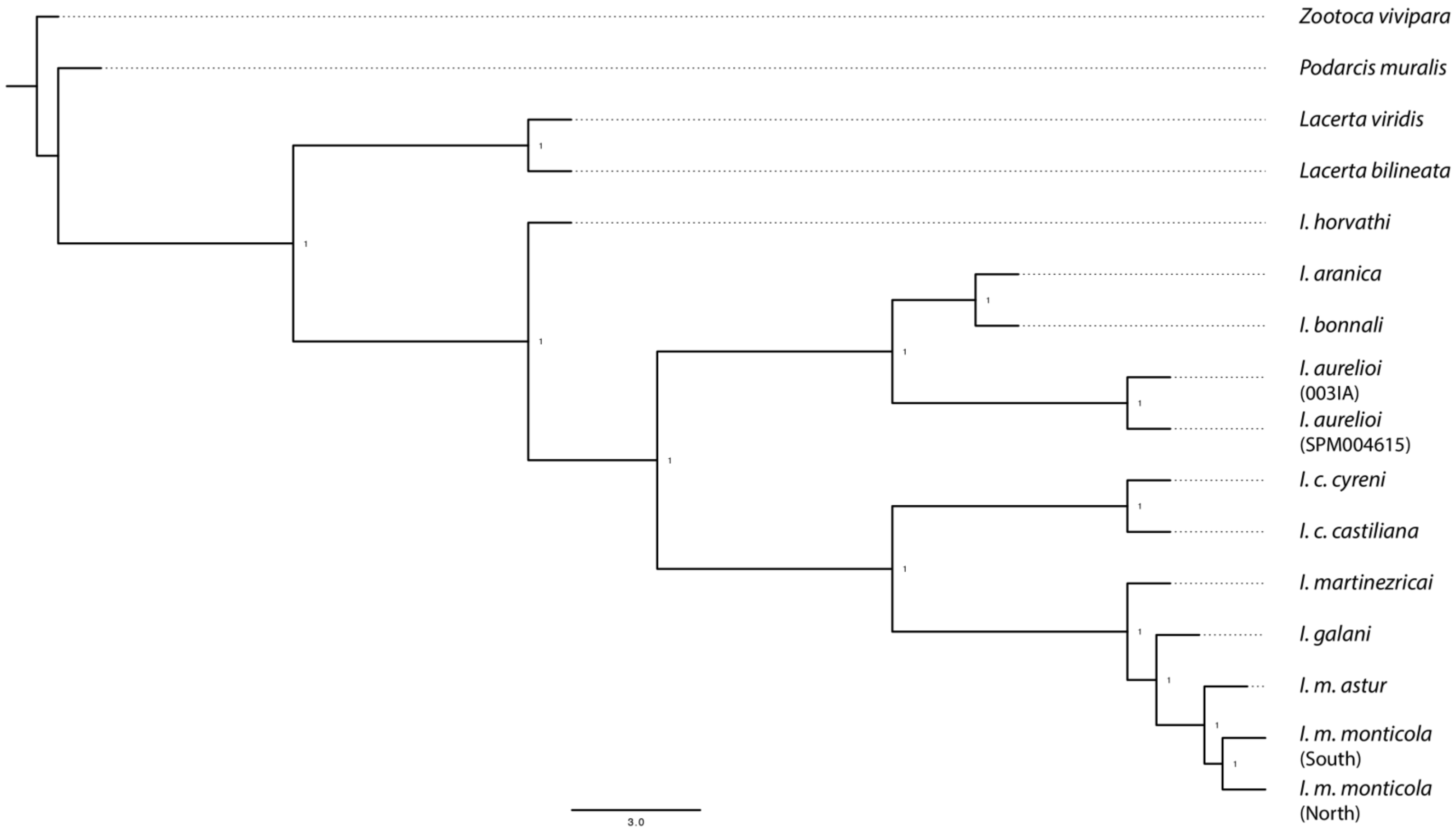


**Fig. S2. Maximum Likelihood consensus tree of the rock lizards of the genus *Iberolacerta*, including some other Lacertidae genera as outgroups.** A dataset containing >67M SNPs was divided in 100-kb non-overlapping windows across the autosomes to run IQtree v2 with GTR-ASC model and 1,000 bootstrap replicates. Best trees were combined with ASTRAL-III into a consensus tree, whose topology is concordant with Bayesian species tree. All nodes have maximum support.

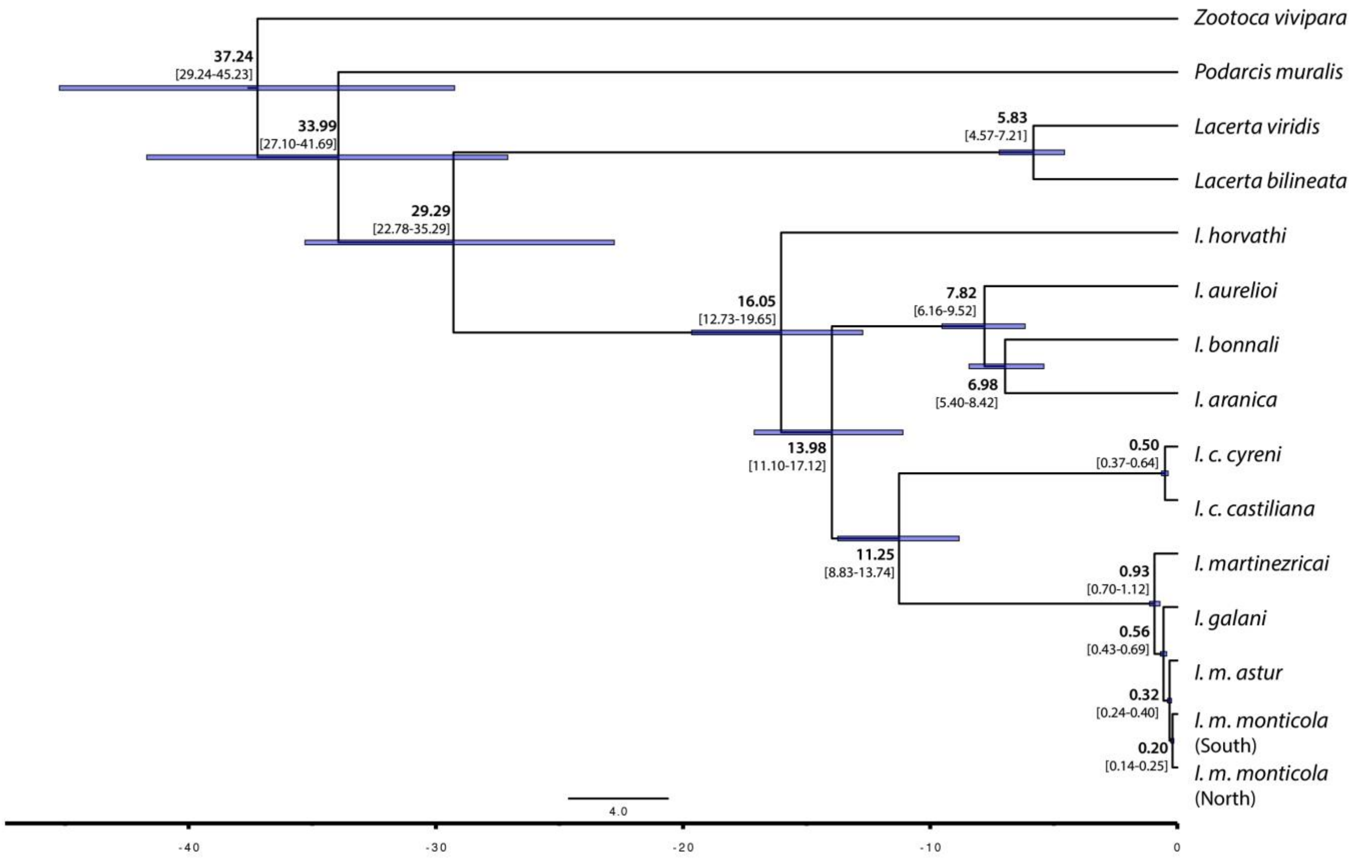


**Fig. S3. Time-calibrated Bayesian species tree, performed with SNAPP implemented on BEAST2.** The analysis was based on a dataset comprising over 123,000 unlinked SNPs (uSNPs). The crown age of the phylogeny was constrained and dated according to García-Porta et al. (2019). All nodes show posterior probabilities of 1. Mean divergence times are indicated in bold, with the corresponding 95% highest posterior density (HPD) intervals shown in square brackets.
